# DECODE-ing sparsity patterns in single-cell RNA-seq

**DOI:** 10.1101/241646

**Authors:** Shahin Mohammadi, Jose Davila-Velderrain, Manolis Kellis, Ananth Grama

## Abstract

An inherent challenge in interpreting single-cell transcriptomic data is the high frequency of zero values. This phenomenon has been attributed to both biological and technical sources, although the extent of the contribution of each remains unclear. Here, we show that the underlying gene presence/absence sparsity patterns are by themselves highly informative. We develop an algorithm, called DECODE, to assess the extent of joint presence/absence of genes across different cells, and to infer a gene dependency network. We show that this network captures biologically-meaningful pathways, cell-type specific modules, and connectivity patterns characteristic of complex networks. We develop a model that uses this network to discriminate biological vs. technical zeros, by exploiting each gene’s local network neighborhood. For inferred non-biological zeros, we build a predictive model that imputes the missing value of each gene based on activity patterns of its most informative neighbors. We show that our framework accurately infers gene-gene functional dependencies, pinpoints technical zeros, and predicts biologically-meaningful missing values in three diverse datasets.

---

Single-cell technologies are revolutionizing the depth that we can investigate cell-type/tissue heterogeneity in health and disease. Among these technologies, single-cell RNA-seq (scRNA-seq) aims to provide a means to quantify this heterogeneity across individual cells at the transcriptome level. This task, however, is challenging due to both biological and technical reasons, including the intrinsically stochastic nature and spatiotemporal dependency of transcription, as well as the challenge of comprehensive sampling of transcript reads within single cells ^1, 2, 3^. These factors jointly influence the structure of single-cell transcriptomic profiles and result in sparse gene expression matrices with an inflated number of zero values, even for genes that show high expression in bulk samples ^4^. To overcome this difficulties, new methods able to distinguish biological vs. technical sources of such sparsity, and to predict the expression value of undetected genes, are needed ^2^.

Here we show that, rather than only a limitation, transcriptome sparsity provides information that reflects biologically meaningful associations. We present a novel framework (DECODE) that enables accurate missing value imputation by network-based modeling of scRNA-seq sparsity patterns. We capture the inherent structure embedded in the sparsity patterns by inferring a gene-gene co-dependency network, where genes are linked based on the statistical dependency of pairwise presence/absence profiles across cells. We demonstrate that this new type of network relates functionally-linked genes, uncovers gene modules that are highly enriched for cell-type specific functions, thus providing meaningful biological information. We then exploit the network structure to predict the cell-specific activity for each gene. Specifically, we quantify the probability of “functional” activity for each gene in each cell, based on aggregate evidence from its first-degree neighbors, adjusting the information flow adaptively for each gene-cell pair.

Our framework provides several predictions regarding the context-specific activity of genes and their functional modularity. We test the accuracy and biological relevance of the predictions using three independent and diverse datasets, encompassing different technologies and heterogeneous tissues/cell types, finding strong support in each case.

Finally, we use DECODE to analyze Type 2 diabetes (T2D) in the context of pancreas single-cell transcriptomics as a case study. Interestingly, by straightforward application of DECODE, we identify cell-specific and T2D-specific pathways supported by functional networks, protein-protein interactions, and distinct gene expression patterns. Our results demonstrate that our framework successfully exploits sparsity patterns in single-cell data to impute missing values and to enable data integration for the study of human biology and disease.

## Deciphering the sparsity structure of single-cell transcriptomic profiles

To detect potential biological signals in the sparsity structure stemming from zeros in single-cell transcriptional profiles, we introduce a new measure, the gene co-dependency index (CDI), that quantifies the degree of deviation of the observed joint binary activity profiles of gene pairs from an independent, null model. For each pair of genes, in an information-theoretic framework, CDI quantifies the information provided by a non-zero observation for one gene about the probability of a non-zero observation for another gene. Formally, CDI quantifies the deviation of the joint gene co-occurrence frequency from that expected given the gene marginal probabilities (see Methods). The pairwise computation of the CDI results in a weighted, undirected network connecting all pairs of genes based on the degree of dependency in their activity profiles.

Using the CDI, we built gene co-dependency networks for three separate datasets, spanning different cell types, technologies, and species (Supplementary file 1). In order to analyze the structure of the networks, we considered gene-gene edges above a CDI threshold value of 5 (P-value *≤* 10^−5^). We studied several topological properties of the resulting networks, including the clustering coefficient, average path length, modularity, density, and maximum connected component size (Supplementary Table S1). We also studied how these properties changed with varying CDI threshold values (Supplementary Figure S2), and found that each of these properties was stable and changed smoothly up to the CDI=5 threshold, though some became unstable for more stringent CDI thresholds given the increasing sparsity resulting from highly-stringent threshold.

The resulting networks for human pancreas, human PBMC, and mouse brain single-cell data displayed several properties characteristic of real-world complex networks in terms of their clustering coefficient, degree distribution, connectivity preference patterns, and modularity. Their degree distribution showed an approximately linear decay, in the log-frequency as a function of log-degree, which indicates a heterogeneous distribution characterized by a majority of genes having low connectivity and a small group of highly connected genes or hubs (Figure 2a top). In addition, genes with higher degrees connected with genes with similarly high degree, while genes of lower degree did not show a clear preference (Supplementary Figure S1). Although the networks showed qualitatively similar properties, the mouse brain network deviated the most, showing higher density and more highly connected genes.

**Figure 1:**
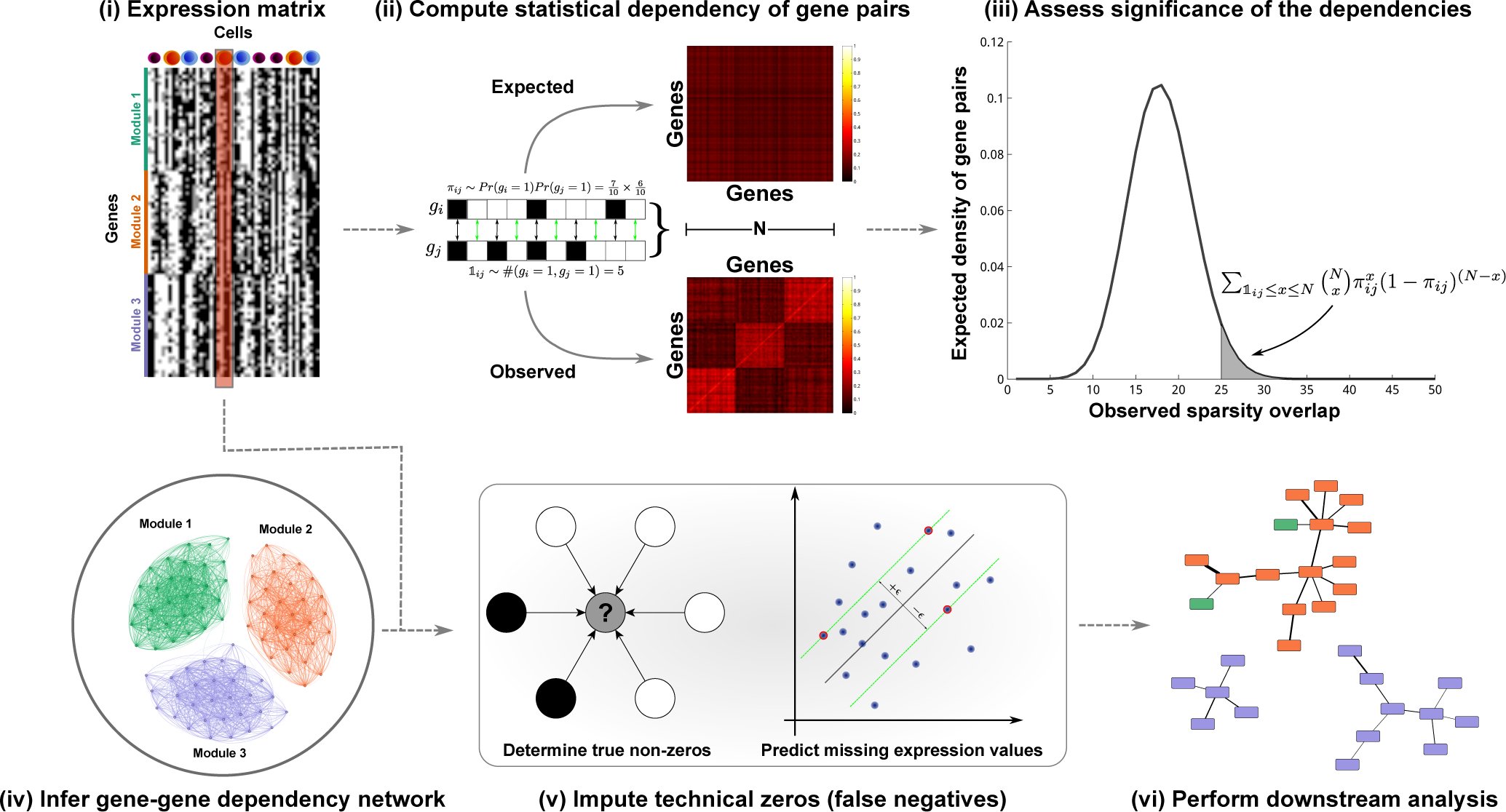
Summary of our approach.

**Figure 2:**
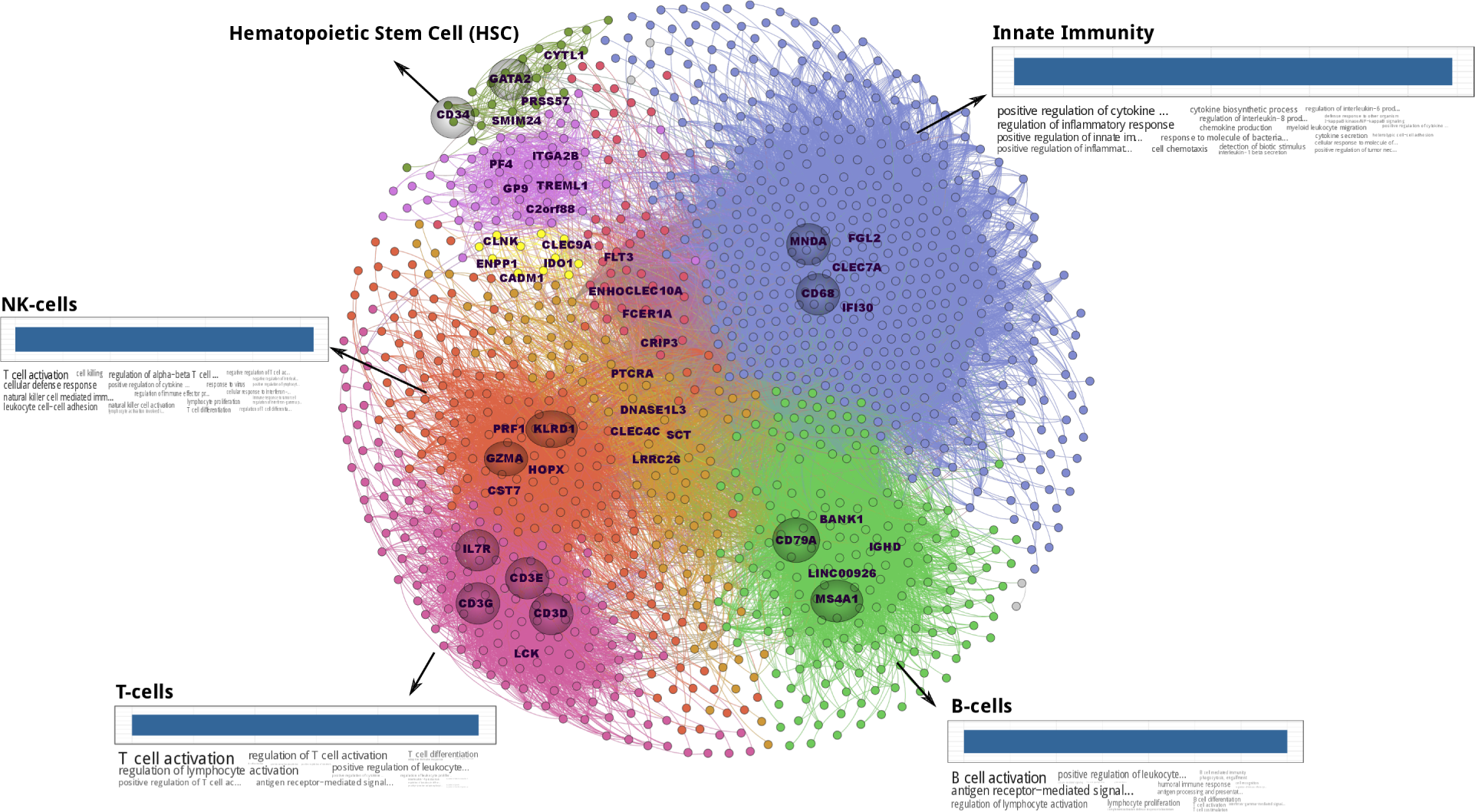
Characterization of gene-gene co-dependency network. Co-dependency network of the PBMC dataset. Each module is color coded accordingly and top four largest modules are annotated with their enriched functions. These modules coincide with the underlying cell types in the PBMC dataset. Top-5 high-degree nodes within each module are marker. Many of these genes (marked with shaded circle) are well-established cell type markers.

We then sought to test whether the resulting gene-gene networks include gene modules involved in shared functional processes and distinctive of cellular states of cell subpopulations. We used a modularity maximization algorithm^5^ to identify gene modules in each dataset (Supplementary file 2). The gene network in the mouse brain dataset partitioned into 5 gene clusters, of which 4 correspond to cell type-specific genes and pathways related to the underlying cell types in the population, including neuronal, microglial, astrocyte, and oligodendrocyte genes. Genes with glial functions showed higher interconnectivity among each other and higher separation from genes with neural functions. The pancreatic gene network partitioned into five modules, of which the top four mapped to endocrine genes (active in alpha and beta islets) and exocrine genes (active in acinar and ductal celltypes). The PBMC gene network partitioned into 10 modules, the largest of which corresponds to genes active in monocytes and dendritic cells. The remaining genes in this last network formed distinct modules that closely match functions associated with the cell-types in the underlying population. Notably, we identified a gene module corresponding to genes active in hematopoietic stem cells (HSCs), suggesting that HSCs are represented in the dataset, although these were missed during the annotation phase of the original data.

We next examined the co-dependency network inferred from PBMC single-cell data in greater detail. We first visualized the network and color-coded the modules according to their neighborhoods (Figure 2b). Experimentally, the PBMC dataset was annotated to contain four cell types: NK-cells, T-cells, B-cells, and myeloid-derived cells (monocytes and dendritic cells). Indeed, the gene modules recovered in the codependency network correspond to genes with known functions in each of the four cell types, and these functions are significantly enriched in each case. In addition, we studied the most connected (hub) genes within each module, which, in our case, represent those genes having high co-dependence with other genes. We found that many of these hubs correspond to known canonical marker genes commonly used to characterize and isolate the corresponding cell types, even though they were inferred entirely *de novo* from the collective sparsity pattern associations. This further indicates that the introduced co-dependency network indeed recovers relevant, functional modularity. In addition to the four originally annotated cell types, we found within the discovered HSCs module the canonical marker *CD34* as a hub, further suggesting the presence of a subpopulation of cells with a HSC state.

We next compared the introduced CDI measure with two commonly-used measures of gene functional interaction: correlation, and mutual information (Figure S4). We focused on the subspace of marker genes and compared the ability of each method to recover pairs of functionally-related genes, defined as pairs of genes that are significantly co-expressed in a given cell-type. In the MouseBrain and Pancreas dataset, the CDI had the highest AUC (0.91 and 0.89), whereas in the PBMC dataset it falls on the second place with AUC of 0.85. Interestingly, gene-gene expression correlation had the worst performance in all three cases, and its performance worsens as sparsity increases (e.g., in Pancreas and PBMC datasets).

## Inferring gene activities using gene-gene dependency network

We next sought to utilize the gene co-dependency network to infer whether or not a given gene is functionally active, in other words, to distinguish between the biological versus technical zeros in a single-cell dataset. Following the principle of *guilt-by-association*^6^, we reasoned that the activity of a given gene can be collectively inferred from that of its neighbors, which are likely to be involved in common processes.

Given a gene *g*, we infer its functional activity from an optimally-informative neighborhood around it, which is constructed individually for each cell *c*. More specifically, we sort all gene neighbors based on their co-dependency *p*-values, label them as either expressed or not in *c*, and find the optimal number of top-ranked genes that are maximally enriched for expressed genes. To this end, we compute the tail of a hypergeometric distribution for neighborhoods of different sizes. After finding the optimal size, we use a dynamic programming approach to compute the exact minimum hypergeometric (mHG) *p*-value ^20^ for the set of neighboring genes. This method results in a matrix of *p*-values for functional activity for every gene *g* in every cell *c*, which we use to discriminate technical and biological zeros.

As a benchmark, we used cell type-specific markers, given their highly selective expression domain. Qualitatively, the expression matrix of markers/cell types exhibits a block-diagonal structure (Figure 3). However, these blocks are highly sparse, due to the nature of single-cell expression profiles. Applying DECODE, we observed that it fills in the missing values among block-diagonal entries, thus reducing false negatives, while leaving off-diagonal entries as zeros, indicating that it does not inflate true negative values.

**Figure 3:**
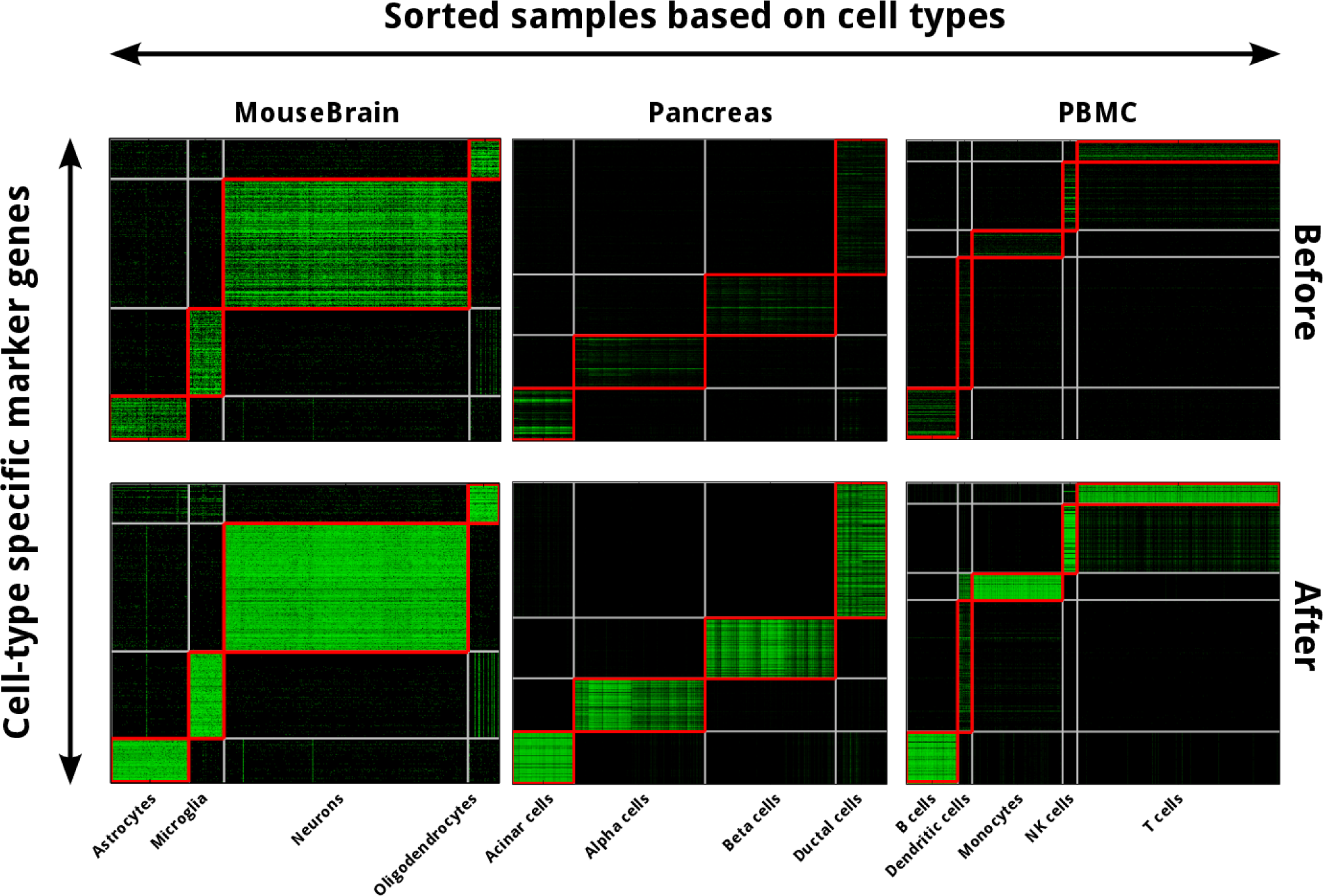
Illustration of DECODE-ing marker subspace within different datasets. Distribution of nonzeros in the original expression matrix, restricted to the subspace of marker genes, and after applying DECODE. Block diagonal elements, marked with red rectangles, show matching markers/cell types. Offdiagonal entries are treated as true biological zeros, or true negatives.

To quantitatively validate our method, we next compared its performance to an alternate approach, *scImpute*^7^. We treated block diagonal elements as true-positives and off-diagonal as true-negatives and computed sensitivity and specificity for each method. In all cases, our network-based approach outperforms *scImpute*, with an average of 20.4%and 29.4%increase in sensitivity and specificity, respectively (Figure 4a). However, depending on the sparsity of the dataset, the difference in performance is biased in one or the other direction of sparsity. For dense datasets (e.g., MouseBrain), the difference is mainly observed in higher specificity of the predictions; whereas in the case of sparse datasets, the difference is mainly observed in sensitivity. In other words, if the majority of non-zero markers were known, our approach does not attempt to falsely declare additional non-zeros. At the same time, if non-zeros are sparse, our method accurately identifies them.

**Figure 4:**
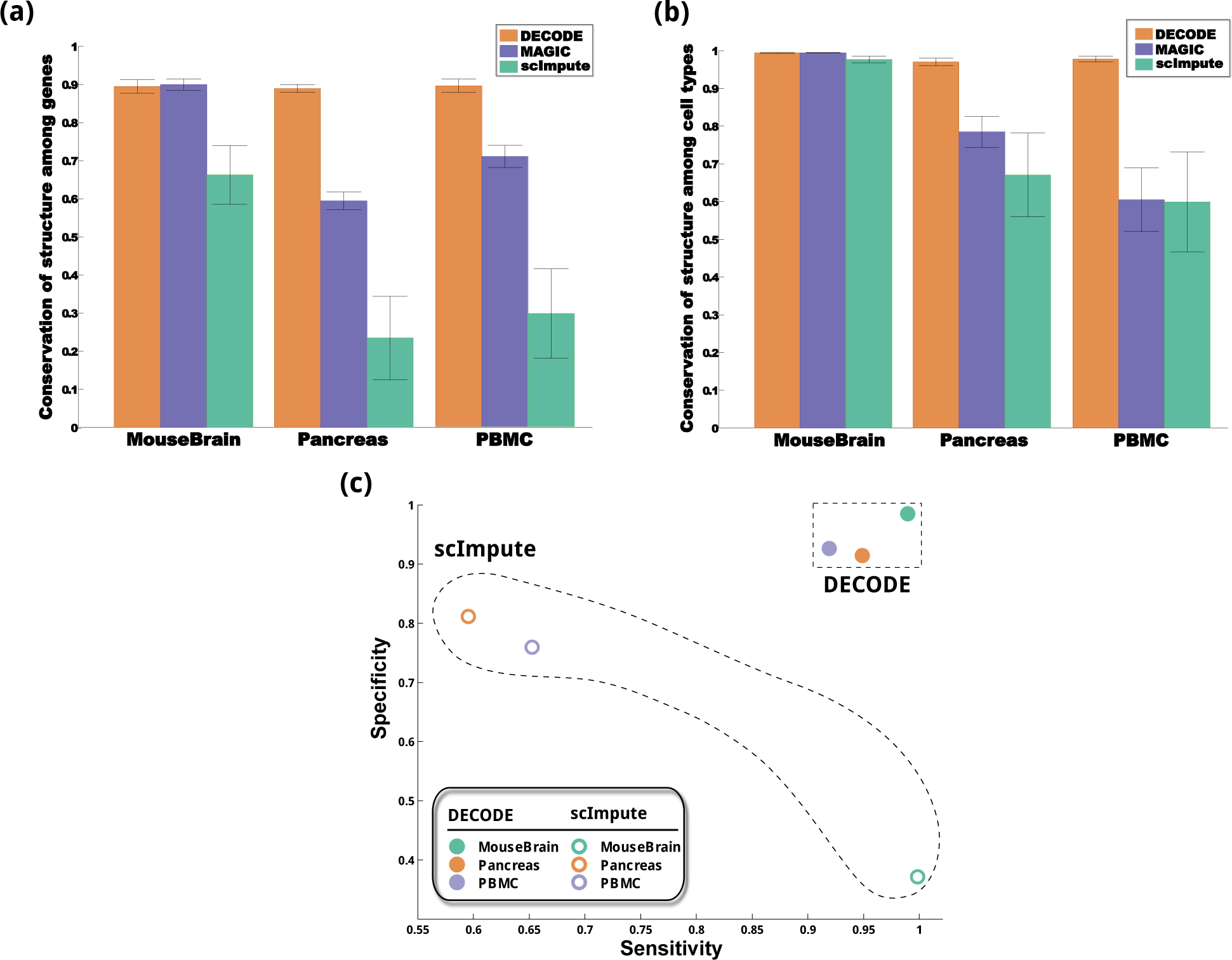
Performance evaluation of DECODE method in imputing missing values. (a) Sensitivity and specificity of DECODE versus scImpute. Our method outperforms scImpute in terms of both measure in all datasets. (b) and (c) Cross-validation missing value imputation using 5×5 fold cross-validation. We randomly permuted rows and columns of the expression matrix and split each dimension into 5 equally-size sections. Then, we masked each block at a time and predicted its values using the rest of matrix with different methods. Finally, we assessed the ability of predictions from each method to reconstruct the underlying correlation structure between genes and cells.

## Supervised learning of technical zeros using gene dependency network

We next sought to infer the expression values for genes that are predicted to be false negatives (technical zeros). To this end, we use a predictive model based on support-vector regression – *∈-SVR*. For each gene in each given sample, we extract the members of its informative neighborhood and use them as features for training. We partition samples into two groups based on the missing pattern of the given gene. We use the group with nonzero values to train the regression model, which we use to impute the technical zeros in the other group (with missing values). We fine-tune SVR parameters using a cross-validation approach (Supplementary Figure S6). Overall, we observed better performance using a linear kernel, which we used throughout the study.

We evaluate the predictions using 5-fold cross-validation for both genes and cells (combined 25fold). We randomly split the genes (rows) and cells (columns) of the expression matrix into 25 gene-cell groups. We then iteratively zeroed the expression values for all genes in one of the 25 blocks, and then used DECODE to predict their expression values. We evaluated imputation quality based on the ability to recover genes of similar function and cells of similar type through their correlation values. Specifically, we first used the original observed data to compute the true correlation of all gene pairs and all cell pairs for the block being evaluated. We then zeroed all expression values for that block (for all genes in the block in the cells in the block), ran the DECODE algorithm on the newly-constructed matrix with 24 observed blocks and 1 zeroed-out block, and re-computed the within-block correlation for all gene pairs and all cell pairs. As this part of the evaluation focused on the ability to infer the true expression values of genes that were previously determined to be non-zero, we only evaluated this correlation for the subset of genes that are predicted to be non-zero.

We performed the same procedure for *DECODE* and two additional imputation methods, *scImpute* and *MAGIC*, and compared their performance (Figures 4b and 4c). For dense datasets (e.g. MouseBrain), we found that all methods are able to recover cell-to-cell relationships, and both *MAGIC* and *DECODE* were able to recover gene-gene relationships with high-quality. For sparse datasets, however, only DECODE was able to faithfully recover both gene-gene and cell-cell relationships.

## Application to Type 2 Diabetes (T2D)

We next sought to study the impact of our imputation approach on elucidating disease-related gene modules in specific cell-types. We focused on Type 2 diabetes, a chronic metabolic disorder characterized by altered glucose processing resulting from pancreatic islet damage. We first constructed a gene-correlation network using the DECODE-imputed transcriptional profile of human pancreatic cells. We then incorporated a set of cell type-specific genes that were recently found to be differentially expressed in T2D patients relative to healthy controls ^8^. Specifically, we extracted from the global pancreas gene correlation network, the subnetwork produced by the vertex-induced extraction of the T2D differential genes. We found that the distribution of correlation scores exhibits a clear separation near 0.2, which we used as a threshold value to binarize the network (Supplementary Figure S7). We also constructed for the same genes a physical interactome using protein-protein interactions obtained from InWeb IM^9^. Finally, we used the binarized gene correlation network as a mask to select context-specific protein-protein interactions involved in cell type-specific mediation of T2D pathology. We identified three different gene modules that are both differentially expressed in T2D and supported by physical protein interactions (Figure 5).

**Figure 5:**
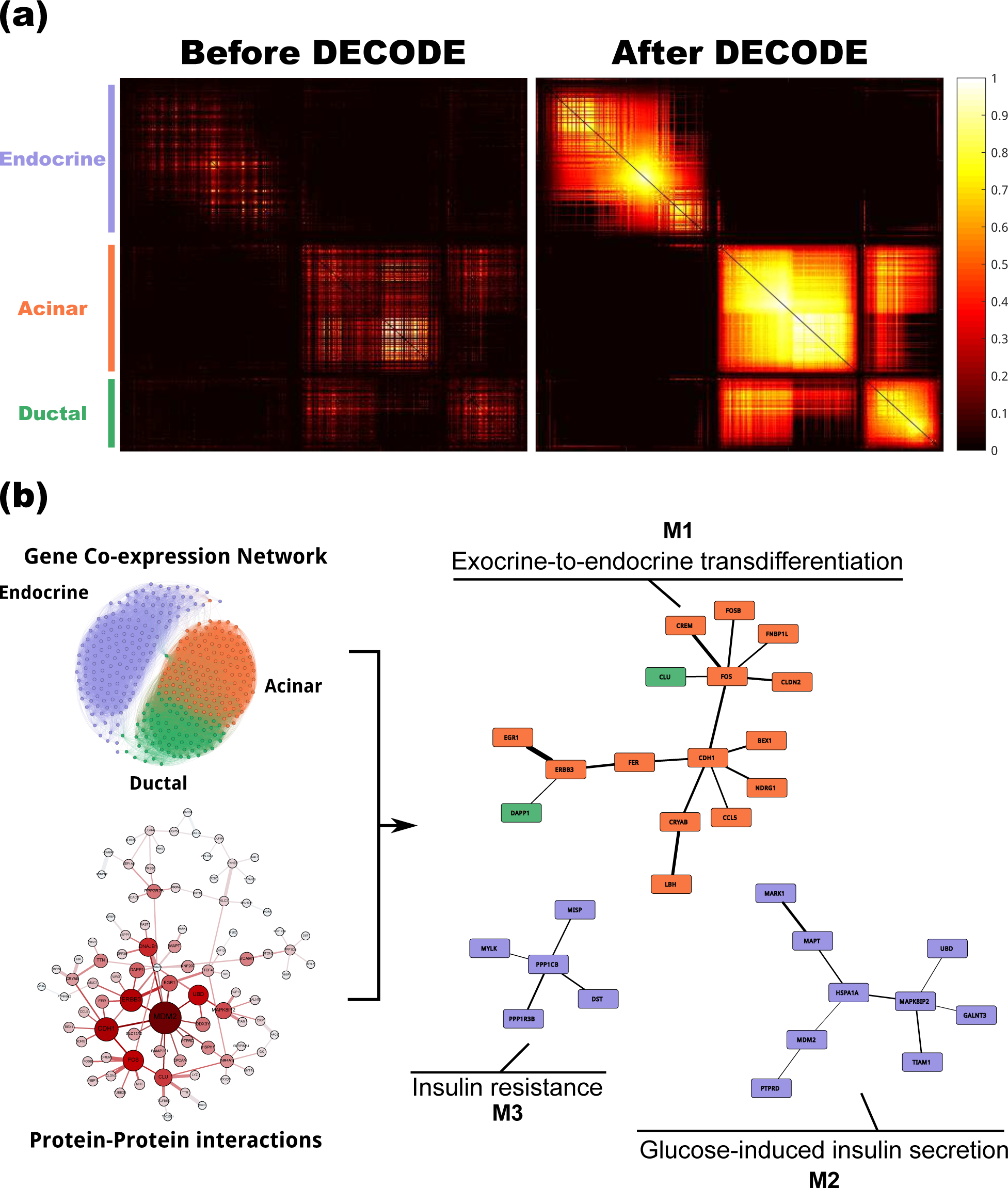
Combining Gene Co-expression Network and Protein-Protein Interactions to identify T2Dspecific pathways. (a) Gene co-expression network among T2D-associated/cell type-specific gene, constructed using the original expression profile and the one imputed using DECODE framework. (b) Combining the co-expression network computed from imputed profile with the protein-protein interactions (PPIs) downloaded from the InWeb IM database. We used co-expressed genes as a filter to select for active/context-specific PPIs. We identify three different pathways that are directly related to insulin signaling, secretion, and resistance.

Module *M1* includes key elements previously identified as key regulators of a trans-differentiation process from acinar to insulin secreting *β*-cells^10^. Cadherin-1 (*CDH1*), the central hub in this module, is a calcium-dependent cell-cell adhesion glycoprotein that is necessary for in-vivo pancreatic exocrine-to-endocrine cell transdifferentiation ^10^. Consistently, two other genes in the module, *CLDN2* and *NDRG1*, are also associated with cell-to-cell adhesion. This transdifferentiation process is also dependent on the phosphatidylinositol 3-kinase (PI3K) signaling pathway. Interestingly, we found an additional gene (*FER*) within the same module, which encodes a tyrosine-protein kinase that is involved in the insulin receptor signaling pathway via PI3K. From a different point of view, this transdifferentiation module also includes the early growth response gene (EGR-1), a transcription factor that is rapidly induced in response to glucose in *β*cells. This rapid expression change is directly linked to insulin secretion and is also elicited by cyclic AMP (cAMP) agonists^11^. Finally, three additional genes in the module (*FOS, FOSB*, and *CREM*) mediate cAMP response. Collectively, these results are consistent with molecular mechanisms known to have a role in the transdifferentiation of exocrine to endocrine cells. On the other hand, its discovery through T2D differential expression, suggest a potential role in disease etiology. The genes in the other two discovered modules are also associated with key processes underlying T2D biology. Module *M2* is involved in glucose-stimulated insulin secretion (GSIS) dependent on MAPK (e.g., TIAM1 and Rac1) ^12^, whereas module *M3* contains genes that are annotated with insulin resistance (*PPP1CB* and *PPP1R3B*). Overall, these results demonstrate the utility of the information gained through network-based transfer of cell-specific gene activity for imputation.

## Discussion

Here we presented a novel approach for modeling and imputing missing values in single-cell gene expression profiles, which discriminates biological and technical zeros. Our model provides an orthogonal view which differs from previous approaches in that it uses a network of gene-gene interactions instead of cell-cell similarities. In methods based on cell-cell similarity networks, the number of selected neighbors and the choice of diffusion depth has a significant impact on the performance of the method. Choosing a small number may not adequately address the imputation problem, whereas large values can smear the subtle cell state signals. Furthermore, these methods deteriorate the existing biological signal in the sparsity pattern of single-cell expression profile.

The introduced gene-gene co-dependency network differs from previous network types, such as PPI and co-expression networks, and it is solely constructed based on the sparsity pattern of single-cell expression profiles. We showed that the proposed network inference method recovers modules/pathways that are highly cell-type specific within the given cell population without having any prior knowledge as to which subpopulation it is being applied to. Using this network, we imputed missing values in different datasets encompassing diverse platforms, cell types, and species. We expect our framework to of broad applicability, both to construct the gene-gene network and discover cell type-specific modules, and to impute missing values in single-cell gene expression profiles.

## Methods

### Datasets

#### FANTOM 5 gene panel

Focusing on a curated set of condition-specific genes significantly enhances the accuracy of cell type predictions ^13^. Following this logic, a subset of reference genes for mouse and human was defined and used throughout the study. The RLE normalized CAGE peak-based expression tables for human and mouse were downloaded from FANTOM5 ^14^ release (11-Nov-2014), from http://fantom.gsc.riken.jp/5/datafiles/latest/extra/CAGE_peaks. All gene IDs were converted to standard gene symbols using HUGO^15^ and MGI^16^ databases. The final dataset contains 18, 869 genes (1, 073 samples) and 18, 852 genes (1, 829 samples), in mouse and human, respectively. These samples include primary known cell types, tissues, and cell lines. Each gene was preprocessed individually and retained in the feature set if its expression in any of the samples is at least an order of magnitude greater than the median of the rest of cell types. This resulted in 6, 596 genes for human and 4, 875 genes for mouse.

#### Single-cell RNA-seq datasets

1. Mouse Brain^17^: was obtained from NCBI GEO under accession number GSE82187. Among the 1, 208 single striatal cells assayed, 532 were selected that were measured using *microfluidic singlecell RNAseq (Mic-scRNAseq)* technology and span 4 major brain cell types: *neurons, astrocytes, oligodendrocytes*, and *microglial cells*. The original dataset contained 18, 840 genes (24%non-zeros). After preprocessing and limitting to the FANTOM panel, the final data included 2, 334 genes (32.47%non-zeros).
2. PBMC: filtered 4k PBMC dataset was downloaded from the 10x Genomics website (https://support.10xgenomics.com/single-cell-gene-expression/datasets/pbmc4k). Cell annotations were transfered from pure cells in a previous study^18^ by matching to the closest cell. Different subtypes of *T cells* were collapsed. The dataset spans cells involved in both innate and adaptive immunity, including *monocytes* and *dedritic cells*, as well as *T, B*, and *NK* cells. The initial dataset contained 13, 704 genes and 5, 825 cells (9.3%non-zeros). Filtering genes and retaining genes in FANTOM5 gene panel resulted in 2, 398 genes (5.6%non-zeros).
3. Pancreas^19^: processed dataset was downloaded from (https://shenorrlab.github.io/bseqsc/vignettes/pages/data.html)^19^. It measures transcriptomic profile of cells from 3 healthy individuals using inDrop. The four most well-characterized cell types, namely *alpha, beta, ductal* and *acinar cells*, were selected. Cells (a total of 4, 352) span both endocrine and exocrine cells. The dataset contained 17, 434 genes (10.7%non-zeros). The final dataset after preprocessing contains 3, 437 genes (5.8%non-zeros).

### Gene co-dependency index (CDI)

Co-dependency of gene activity was measured by quantifying how informative observing a non-zero for a gene *g*_*i*_ is with respect to observing a non-zero for a gene *g*_*j*_. Assume that *N* represents the total number of cells, 𝕝_*i*_ the total number of observed non-zeros for gene *i* across all cells, and 𝕝_*ij*_ the number of cells in which both gene *i* and gene *j* are jointly active (have non-zero values). Then, *P* (*gi* = 1) will denote the probability of observing a cell with non-zero value for gene *i*, and it is computed as 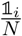. Similarly, the joint probability distribution for a pair of given genes, *g*_*i*_and *g*_*j*_is 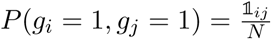 indicating the probability of simultaneously observing both *g*_*i*_and *g*_*j*_to be non-zeros. Assume a null model were genes *g*_*i*_and *g*_*j*_are independent, in which case *P* (*g*_*i*_= 1, *gj* = 1) can be factored as *P* (*g*_*i*_= 1)*P* (*g*_*j*_= 1). The co-dependency of genes is measured based on the deviation of these two terms – i.e., the joint probability from the product of factored marginal distribution. Finally, consider the factorized probability of observing both gene *i* and gene *j* active as *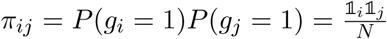*, and assume a binomial distribution as a null model.

Putting the previous pieces together, the co-dependency index (CDI) is defined as follows:

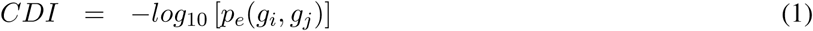

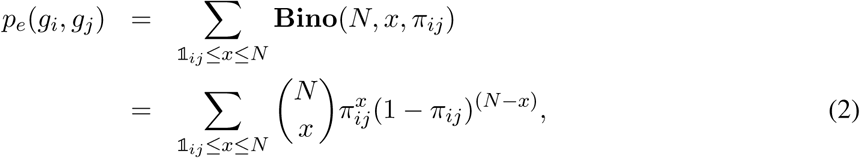

where **Bino**(*N, x, p*) is the probability of observing *x* successes in *N* trials, if the probability of success is *p*.

### Inferring functional activity of genes

Network-based guilt-by-association approach was implemented as follows. A set of significant neighbor genes was selected to act as proxy for gene *i* and to “vote” for its activity status. This set consists of the genes with the most significant co-dependency pattern with gene *i*, not exceeding a total of *𝒦*_*max*_. Define this set by *𝒩* (*i*), and the total number of genes with non-zero values within this set with 𝕝_𝒩_ = ∑_*s∈𝒩*_ (*i*) 𝕝(*s ≠* 0). Denote with 𝕝*k* the total number of non-zeros in cell *k* and let *m* be the the total number of genes. The evidence of activity of proxy genes is aggregated using the hypergeometric tail, testing the null hypothesis:“*H*0: *g*_*i*_= biological zero”, as follows

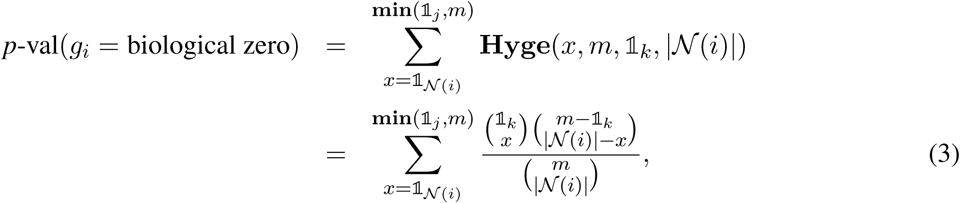

where Hyge(*x, m, k, n*) represents the hypergeometric probability distribution, which calculates the probability of observing *x* white balls when sampling *n* balls from a population of size *m*, and where there are *k* white and *m – k* black balls.

Note that all neighbors of gene *i* are not equally informative and using all |*𝒩* (*i*)| neighbors might not always be optimal. To remedy this problem, multiple neighborhood sizes were tested, computing for each the minimum hypergeometric (mHG) score. The set size minimizing the mHG score was selected. A dynamic programming approach was subsequently used for multiple testing correction, and to assign a *p*-value to the mHG score. This process was originally proposed for geneset enrichment analysis without prespecifying the number of top-ranked genes^20^. In our experiment, a max degree of 100 was chosen to build the network. For computed activity *p*-values, multiple hypothesis testing was corrected using Family-wise Discovery Rate (FDR) adjustment ^21^.

## Computing expression domain of genes and identifying cell type-specific marker genes

For a gene *i* in a cell type *c*, we use Fisher’s exact test to assess the deviation of non-zeros in this context from the background null distribution ofall non-zeros. Let *N*_*c*_ show the total number of cells in cell type *c*, and *p*1 be the probability of observing non-zeros in the background distribution. Thus, the expected number of non-zeros in a cell of size *N*_*c*_ is *p*_1_ *× N_c_*. Positive deviation from this expectation, measured using the tail of the hypergeometric distribution, measures the over-enrichment of non-zeros in that cell type. We use a *p*-value of 0.05 to binarize these memberships. Given a total of *k* cell types, we then assign a binary vector of size *k* to each gene that represents its *expression domain*. Pairs of genes with similar expression domain are used here as standard gold pair of genes that are related in a cell type-specific fashion.

On the other hand, to identify cell type-specific marker genes for cell type *c*, we select all genes whose expression domain has a 1 at position *c*, and zero everywhere else. This provides a list of markers for each of the constituting cell types in the population, which are used to assess sensitivity/specificity of different methods in predicting true negatives in the single-cell expression profile.

## Imputing missing values using *ϵ*-Support Vector Regression (SVR)

Gene missing values were imputed across cells by regressing over the values of the network neighbors of the gene. Support Vector Regression (SVR)(SVR)^22^ is a popular machine learning regression technique, which maximizes the margins around the estimated separator hyperplane from the closest data points on each side. This margin defines an “insensitive” region in which errors are not penalized. In particular, *∈-SVR* defines the insensitive region by employing an *ϵ*-insensitive loss function, in which errors smaller than *ϵ* are ignored, and errors passing *ϵ* are penalized linearly. Formally, *ϵ*-insensitive loss is defined as follows:

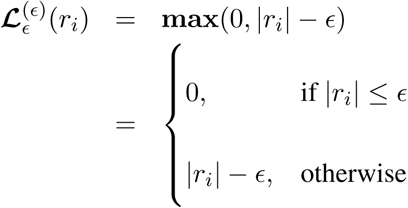

Using this loss function, the primal form of *∈-SVR* can be defined as the following convex optimization problem:

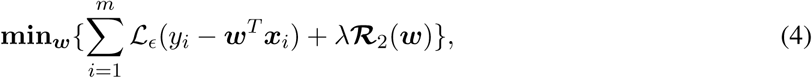

where ***ℛ***_2_ is a *∥ ∥*_2_ regularizer that smooths the weights, while the parameter *λ* balances the importance of fitness versus regularization. This balance is particularly useful in scRNASeq, where the high number of zeros is prone to produce ill-conditioned predictions. To apply our framework, the expression values were first normalized between [0-1] using the 95%percentile. We chose an *ϵ* = 0.01, giving a margin of 1%error. The optimal regularizer (*λ*) was chosen based on cross-validation with exponentially-separated values in the range {2^0^, 2^1^, *…* 2^20^} (Figure S6). *λ* = 2^10^ was found as an optimal trade-off between regularization and minimization of the loss function, and it was used throughout the study.

## Evaluation of Imputed expression values

A 5×5-fold cross-validation experiment was performed in a shuffled expression matrix, splitted across rows and columns in 25 blocks. In each step, one block was masked, and the resulting matrix was used as input to the different methods for prediction. For each block, two matrices, C_*genes*_ and C_*cells*_, were created before and after masking. Similarly, two matrices were created after imputation, denoted by 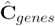 and 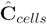, respectively. The conservation of the correlation structure between genes/cells was quantified using a well-established measure for correlation matrix comparison ^23^:

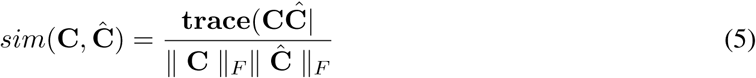

## Availability

Codes and datasets are available to download from http://compbio.mit.edu/DECODE.

## Acknowledgements

This work has been supported through NIH Grants 5R01AI114814-02 and U01-CA198941 and NSF grant CCR-0939370.

## Correspondence

Correspondence and requests for materials should be addressed to S.M. or M.K.

## Authorship

M.K. and A.G. and co-last author on the paper.

## Supplementary Figures

**Figure S1:**
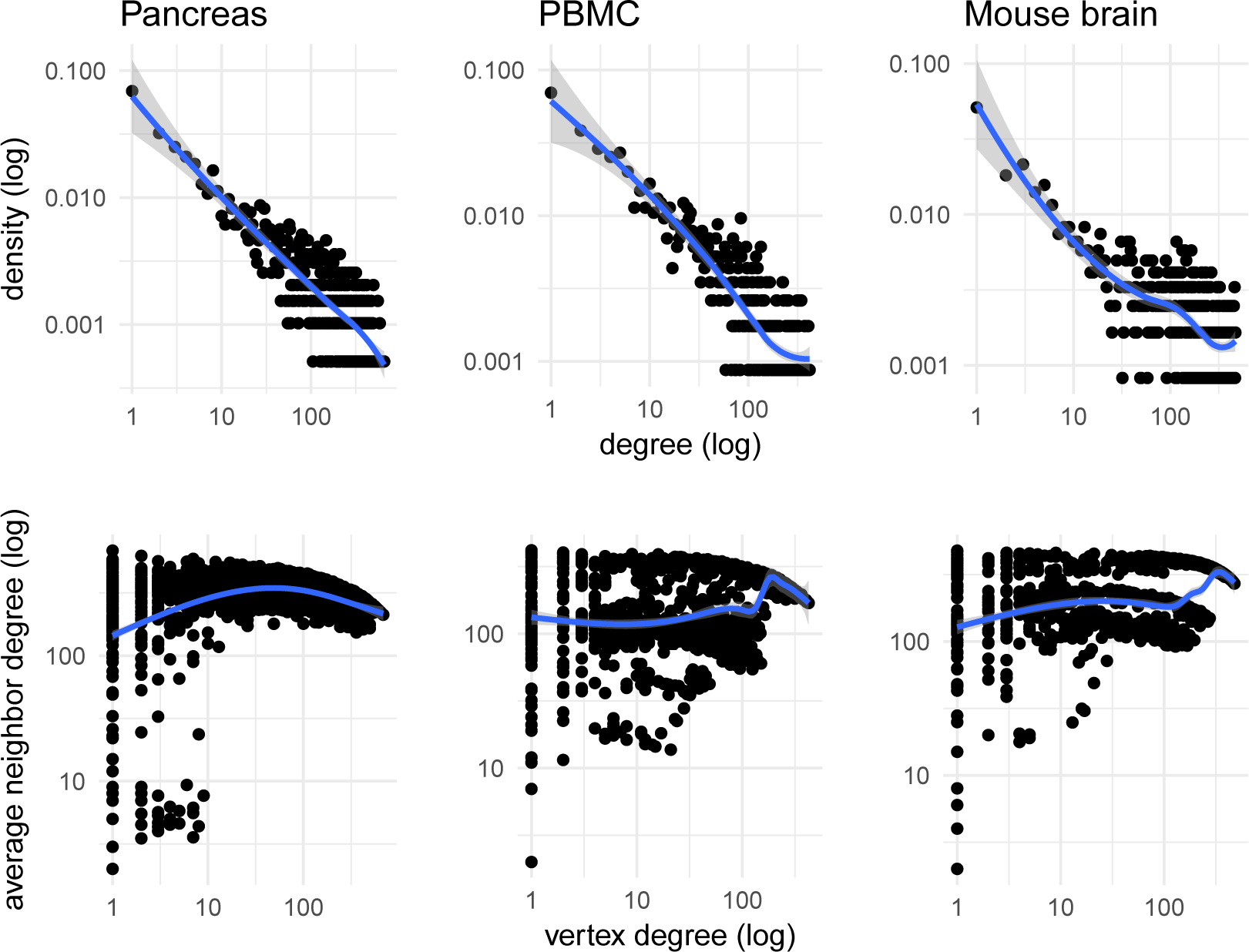
Degree distribution of gene-gene co-dependency network. Gene-gene co-dependency networks resemble many of the common characteristics of complex networks, among which is a log-linear decay in the degree distribution (top panel), and a higher-chance of connectivity between high-degree nodes (lower panel).

**Figure S2:**
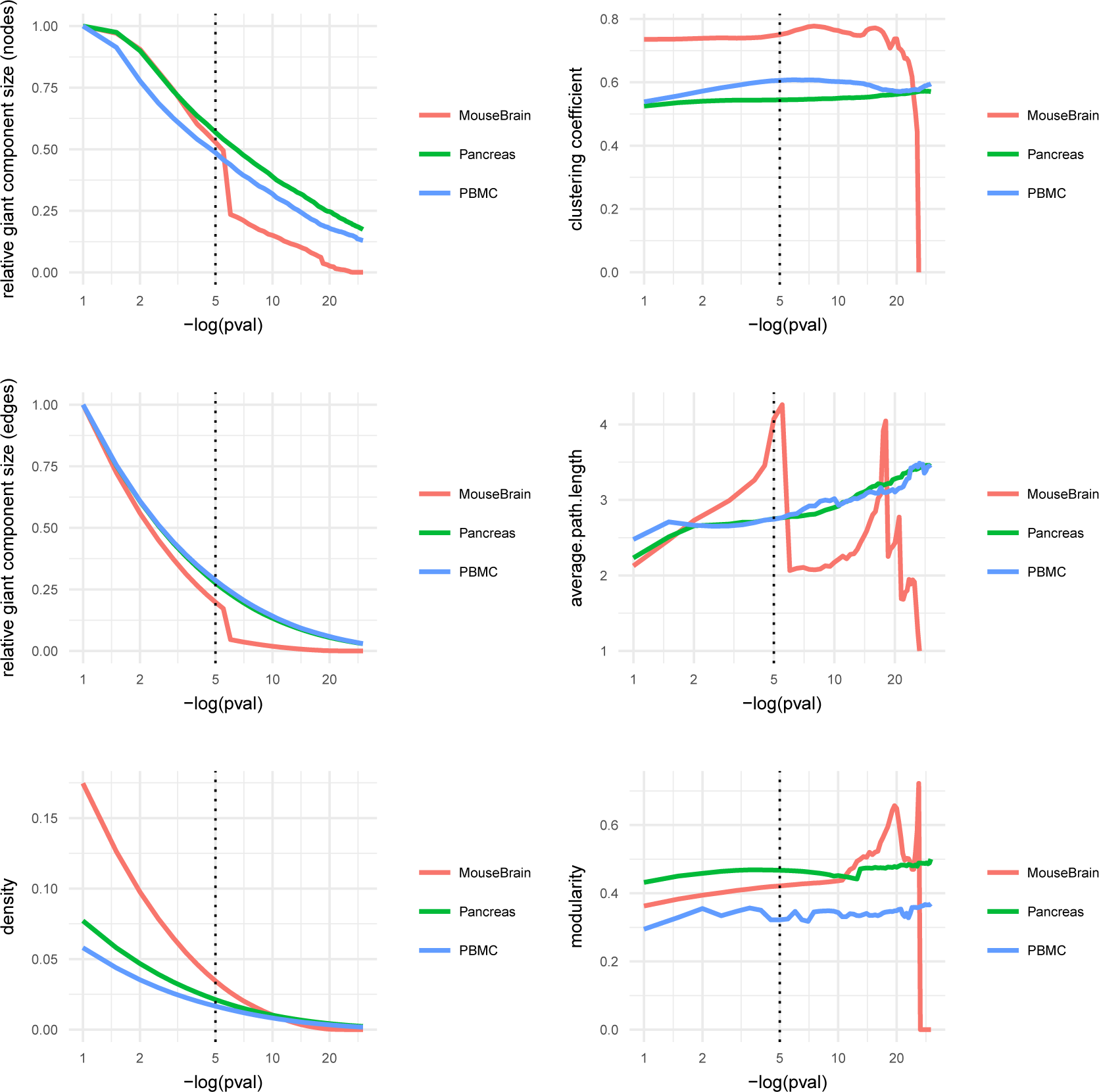
Different network properties as a function of dependency *p*-value. We tried different *p*-value thresholds, ranging from 10^−1^ to 10^−30^ and traced six different network measures characteristics of complex networks as a function of this threshold. We found threshold value of 10^−5^ to be a good cutoff, as the three networks largely converge to a similar size and degree of clustering without displaying abrupt shifts.

**Figure S3:**
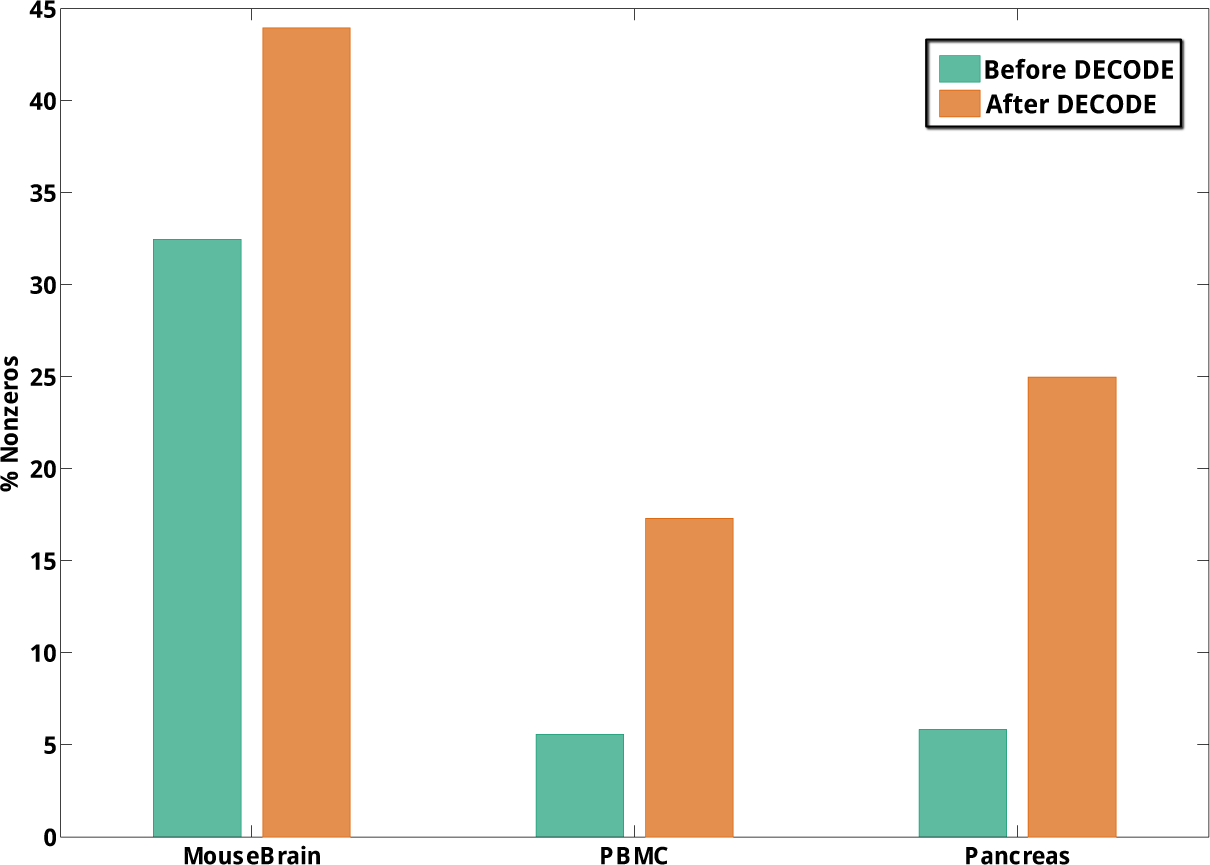
Percent of non-zero values before and after DECODE. For each dataset, percent of imputed non-zeros are different. For the MouseBrain dataset, which is relatively dense to begin with, we observe close to 50%increase in number of non-zeros. For the other two datasets, which are much sparser, we observe 3-5 fold increase in the number of non-zeros.

**Figure S4:**
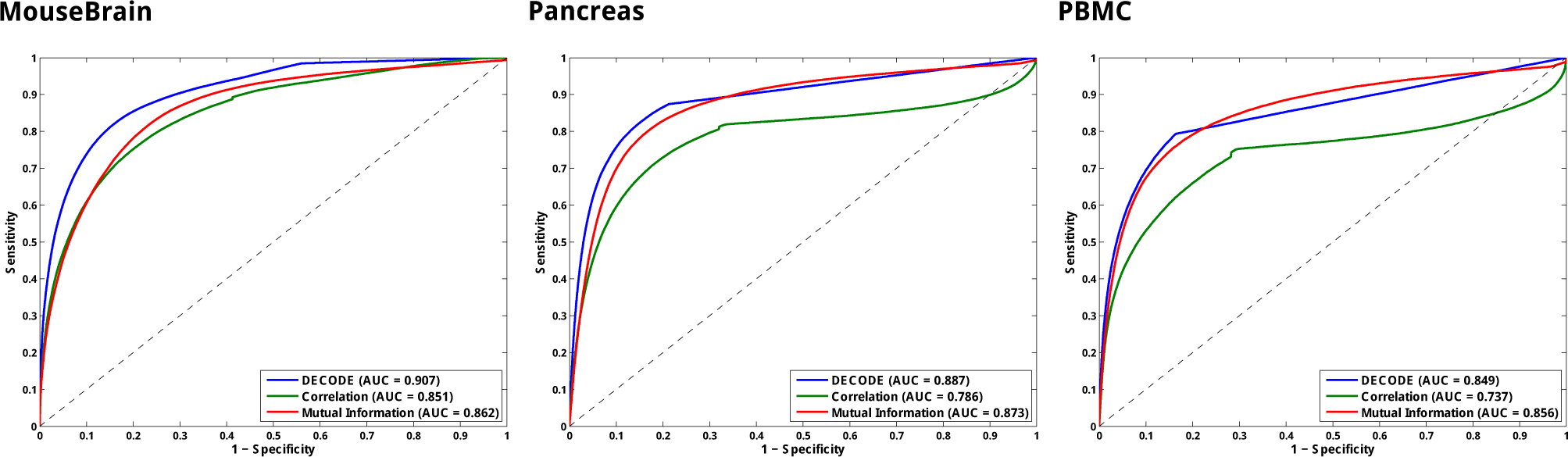
Comparison of CDI versus other measures. Given the known cell types, gene pairs with similar expression domains (expressed in similar cell types) are considered as true-positive set. For each measure, we computed the area under the curve (AUC) of the ROC curve as an indicator of performance. Pairwise gene correlation performs worse in all datasets. DECODE outperforms the other two measures in two out of three datasets, while performs marginally worse in the PBMC dataset. In all datasets, especially Pancreas and PBMC, DECODE plot flattens out, which is due to pairs of gene that are deemed insignificant (with *p*-value 1).

**Figure S5:**
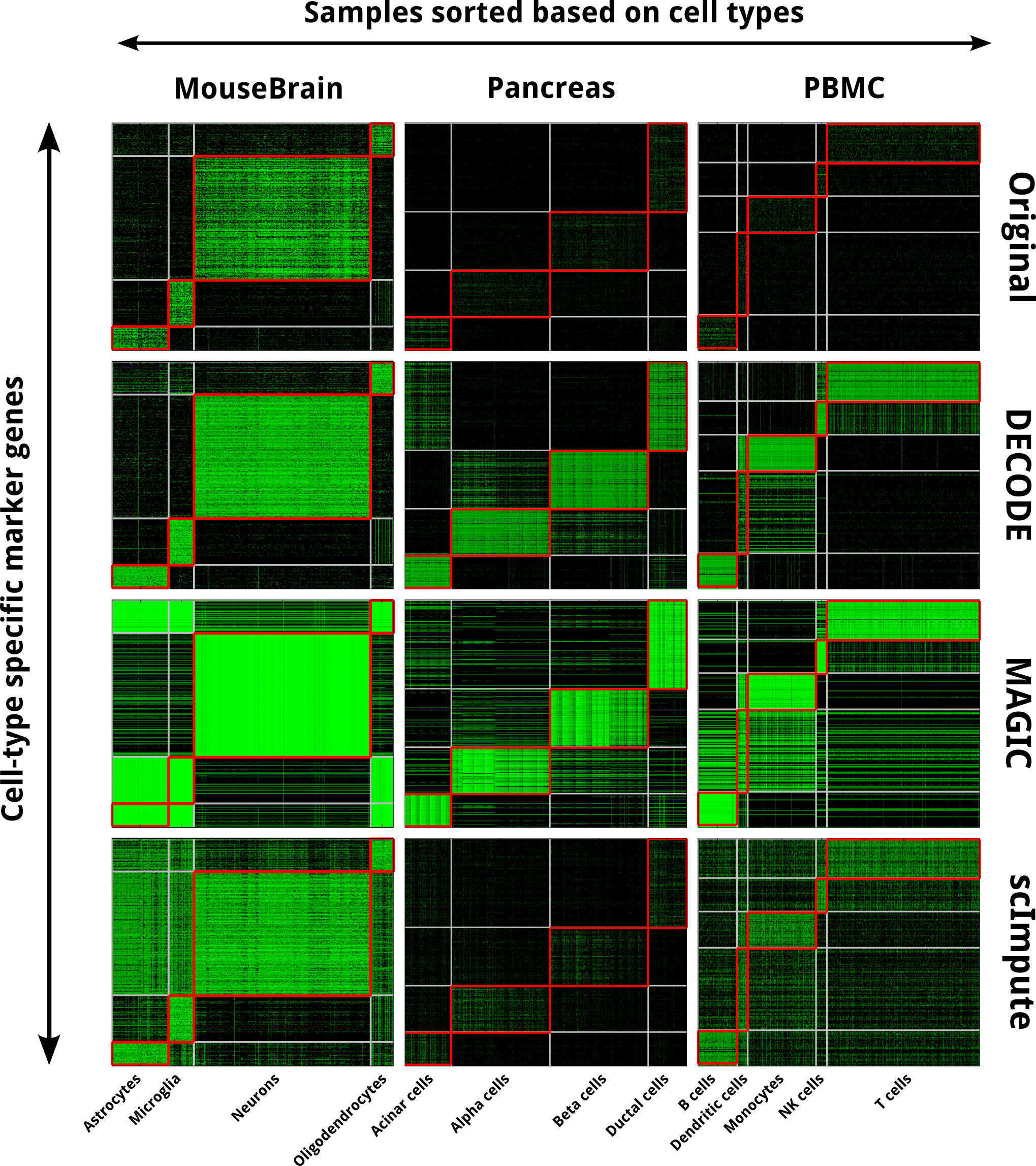
Expression of marker genes in different cell types after imputation. Rows correspond to cell type-specific marker genes and columns are cells, sorted according to their cell types. Ideally, all markers should lie on the diagonal blocks, while off-diagonal elements are treated as true biological zeros. *MAGIC* does not have a mechanism to distinguish between biological versus technical zeros, thus it assigns a value to every zero entities in the expression matrix. *scImpute* only assigns values that it deems to be technical zeros. However, these values are over-estimated in the MouseBrain dataset, while are highly under-estimated in the other two datasets.

**Figure S6:**
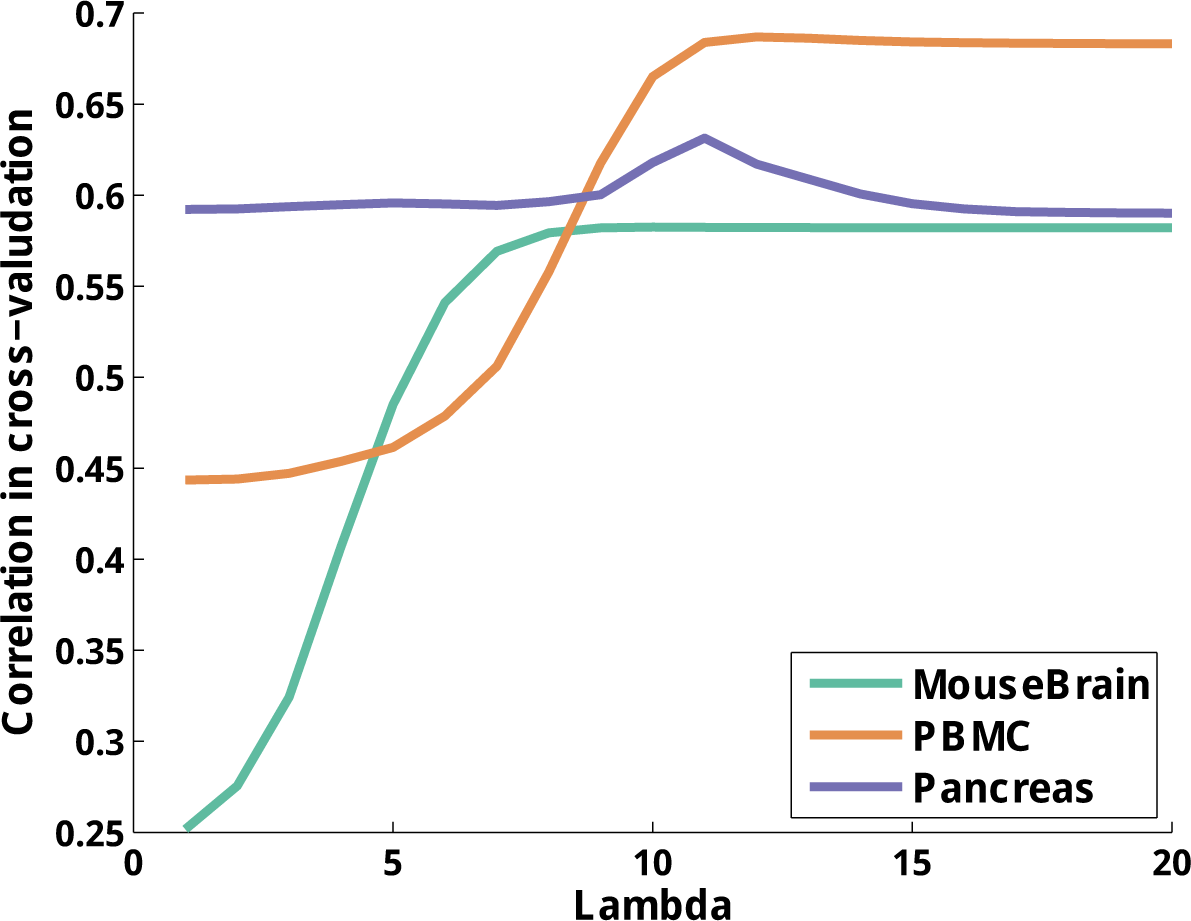
Cross-validation to find optimal regularizer parameter. We tried different values of *λ*, the weight of **ℒ**2-regularization in SVR, and for each case we measured the correlation of predicted values with actual values. X-axes is the *log*2 of lambda. Values of *λ* close to 210 show a good balance of regularization-versus-loss minimization.

**Figure S7:**
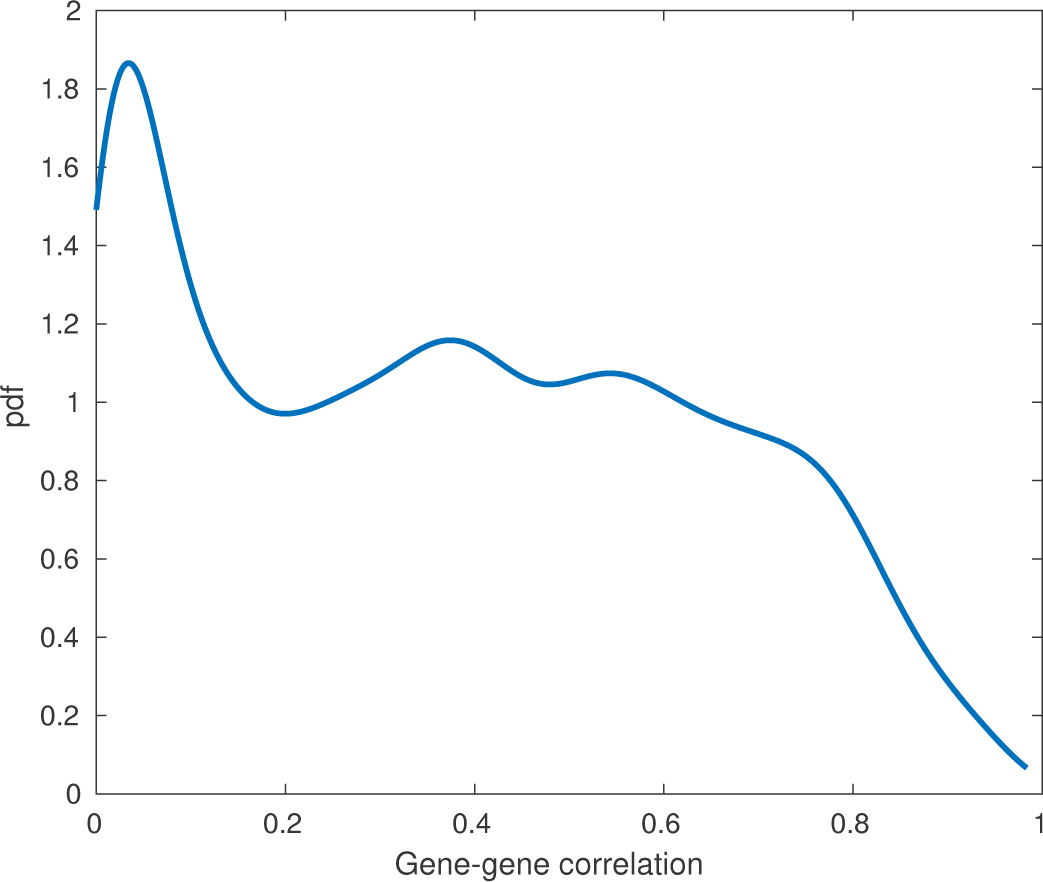
Distribution of gene-correlation scores in the subnet of T2D genes. Distribution of positive correlations has been smoothed using Gaussian kernel and visualized. There is a population shift around 0.2, which we used for thresholding T2D gene-correlation subnetwork.

## Supplementary Tables

**Table S1:** Networks topological statistics. Main topological measures characteristics of complex networks after thresholding dependency networks constructed by DECODE.

| Measures | Networks |  |  |
| --- | --- | --- | --- |
|  | Pancreas | PBMC | Mouse Brain |
| nodes | 1954 | 1146 | 1211 |
| edges | 125605 | 47907 | 94079 |
| mean degree | 128.56 | 83.61 | 155.37 |
| clustering coef. | 0.54 | 0.61 | 0.75 |
| mean shortest path | 2.75 | 2.74 | 4.07 |
| diameter | 10 | 7 | 10 |
| density | 0.07 | 0.07 | 0.13 |

## Supplementary Materials

### Supplementary file 1

This zip file contains co-dependency network between selected genes in each of three datasets in this study

### Supplementary file 2

This zip file contains network modules for co-dependency networks.

### Supplementary file 3

This zip file contains list of cell type specific markers identified in this study.

### Supplementary file 4

This zip file contains predicated gene expression matrices and their corresponding *p*-value matrix for being non-zeros.

